# An early Myc-dependent transcriptional program underlies enhanced macromolecular biosynthesis and cell growth during B-cell activation

**DOI:** 10.1101/561464

**Authors:** Alessandra Tesi, Stefano de Pretis, Mattia Furlan, Marco Filipuzzi, Marco J. Morelli, Adrian Andronache, Mirko Doni, Alessandro Verrecchia, Mattia Pelizzola, Bruno Amati, Arianna Sabò

## Abstract

Upon activation, lymphocytes exit quiescence and undergo substantial increases in cell size, accompanied by activation of energy-producing and anabolic pathways, widespread chromatin decompaction and elevated transcriptional activity. These changes depend upon prior induction of the Myc transcription factor, but how Myc controls them remains unclear. We addressed this issue in primary mouse B-cells, based on conditional deletion of the *c-myc* gene, followed by LPS stimulation. Myc was rapidly induced, became detectable on virtually all active promoters and enhancers, but had no direct impact on global transcriptional activity. Instead, Myc contributed to the swift up- and down-regulation of several hundred genes, including many known regulators of the aforementioned cellular processes. Myc-activated promoters were enriched for E-box consensus motifs, bound Myc at the highest levels and showed enhanced RNA Polymerase II recruitment, the opposite being true at down-regulated loci. Remarkably, the Myc-dependent signature identified in activated B-cells was also enriched in Myc-driven B-cell lymphomas: hence, besides modulation of new cancer-specific programs, the oncogenic action of Myc may largely rely on sustained deregulation of its normal physiological targets.

## Introduction

Mature splenic B-cells can be activated *ex vivo* to re-enter the cell cycle and differentiate into antibody-producing cells, accompanied by massive increases in cell size and RNA content (Kieffer-Kwon, Nimura et al., 2017, Kouzine, Wojtowicz et al., 2013, Nie, Hu et al., 2012, Pogo, Allfrey et al., 1966, Sabò, Kress et al., 2014). This implies a concomitant intensification of the metabolic pathways needed to provide energy and building blocks for macromolecular biosynthesis and cell growth and, in turn, the necessity for the cells to adapt their transcriptional and translational outputs to the augmented cell size and metabolic activity (Marguerat, Schmidt et al., 2012). A key regulator in this overall process is the Myc transcription factor, encoded by the c-*myc* proto-oncogene: indeed, Myc is directly induced by mitogenic signals and, in turn, is thought to orchestrate the plethora of transcriptional changes that foster cell growth and proliferation, as exemplified in cultured mouse fibroblasts (Perna, Faga et al., 2012, Winkles, 1998). In either B or T lymphocytes, c-myc serves as a direct sensor of activating signals (Caro-Maldonado, Wang et al., 2014, Dominguez-Sola, Victora et al., 2012, Kelly, Cochran et al., 1983, Luo, Weisel et al., 2018, Nie et al., 2012, Wang, Dillon et al., 2011) and is required for multiple facets of cellular activation, including metabolic reprogramming, ATP production, ATP-dependent chromatin decompaction, RNA and biomass accumulation, cell growth, etc… (Caro-Maldonado et al., 2014, de Alboran, O’Hagan et al., 2001, De Silva & Klein, 2015, Kieffer-Kwon et al., 2017, Link & Hurlin, 2015, Murn, Mlinaric-Rascan et al., 2009, Nie et al., 2012, Perez-Olivares, Trento et al., 2018, Sabò et al., 2014, Wang et al., 2011). However, how Myc activity impacts on those diverse cellular features remains largely unclear.

Myc binds DNA and activates transcription as a dimer with its partner protein Max (Amati, Dalton et al., 1992, Blackwood & Eisenman, 1991, Kretzner, Blackwood et al., 1992) but its precise contribution to transcriptional programs in cells has been subject of an intense debate in the field in recent years: while multiple studies indicated that Myc can either activate or repress select target genes (Amati et al., 1992, Dang, 2013, Eilers & Eisenman, 2008, Kress, Sabò et al., 2015, Kretzner et al., 1992, Perna et al., 2012), others concluded that it acts instead as a general activator – or *amplifier* – of all expressed genes (Lin, Loven et al., 2012, Nie et al., 2012, Porter, Fisher et al., 2017, Zeid, Lawlor et al., 2018). However, careful scrutiny of the available data (Kieffer-Kwon et al., 2017, Kress, Pellanda et al., 2016, Lin et al., 2012, Lorenzin, Benary et al., 2016, Muhar, Ebert et al., 2018, Nie et al., 2012, Perna et al., 2012, Porter et al., 2017, Sabò et al., 2014, Walz, Lorenzin et al., 2014, Zeid et al., 2018) lends no formal support to this model, suggesting instead that RNA amplification – when present – is one of the consequences of the metabolic and genomic changes that occur during cellular activation and/or transformation (Kress et al., 2016, Sabò & Amati, 2018). Hence, understanding the precise contribution of Myc to global changes in RNA biology will require a fine mapping of direct, Myc-dependent transcriptional programs.

The B-cell system, with its global increase in RNA transcription during cell activation (Pogo et al., 1966), offers a valuable tool to assess the order of events that lead from the triggering of a signaling event to cell cycle entry, cell growth and RNA amplification, and to understand how these events depend upon prior activation of Myc. Toward this aim, we profiled gene expression along with the genomic distribution of Myc and RNA polymerase II (RNAPII) during B-cell activation *in vitro* in wild type and c-*myc* knock-out cells. Our data led to the identification of a specific Myc-dependent transcriptional program occurring within the first few hours upon cell activation, pre-setting the stage for the subsequent global increase in metabolic and biosynthetic activities.

## Results and Discussion

In order to characterize the contribution of Myc to B-cell activation, we took advantage of mice homozygous for a conditional *c-myc* knockout allele (c-*myc*^*f/f*^)(Trumpp, Refaeli et al., 2001). Freshly purified c-*myc*^*f/f*^ and control *c-myc*^*wt/wt*^ B-cells were treated with Cre recombinase, deleting c-*myc*^*f/f*^ with 70-80% efficiency (henceforth c*-myc*^Δ/Δ^) (**Fig. EV1a**) and preventing the LPS-induced accumulation of the *c-myc* mRNA and protein (**Fig. EV1b, c**). Chromatin Immunoprecipitation (ChIP) analysis confirmed rapid binding of Myc to a known target locus (*Ncl*) in *c-myc*^*wt/wt*^ cells (**Fig. EV1d**). *Ncl* and other Myc-dependent mRNAs previously identified in fibroblasts (Perna et al., 2012) responded to LPS in B-cells and required Myc for maximal accumulation, from 4h onward (**Fig. 1a, Fig. EV1e**). While global RNA levels also increased in a Myc-dependent manner (Nie et al., 2012, Sabò et al., 2014), this occurred later (24h, **Fig. EV1f**) concomitant with increases in bulk RNA synthesis and nuclear size (**Fig. 1b**), overall cell size (Kieffer-Kwon et al., 2017, Kouzine et al., 2013, Sabò et al., 2014), as well as S-phase entry (**Fig. EV1g**)(Sabò et al., 2014). As expected (de Alboran et al., 2001, Kieffer-Kwon et al., 2017, Murn et al., 2009, Sabò et al., 2014), all of the above effects were lost in c*-myc*^Δ/Δ^ cells, accompanied by a reduced proliferative response (**Fig. EV1g, h**), residual expansion at 48-72h resulting from the selection of non-deleted c-*myc*^*f/f*^ cells (**Fig. EV1i**). Hence, c-*myc*^*f/f*^ B-cells provide a reliable system to address the role of Myc within the first cell division cycle after LPS stimulation.

**Fig. 1.**
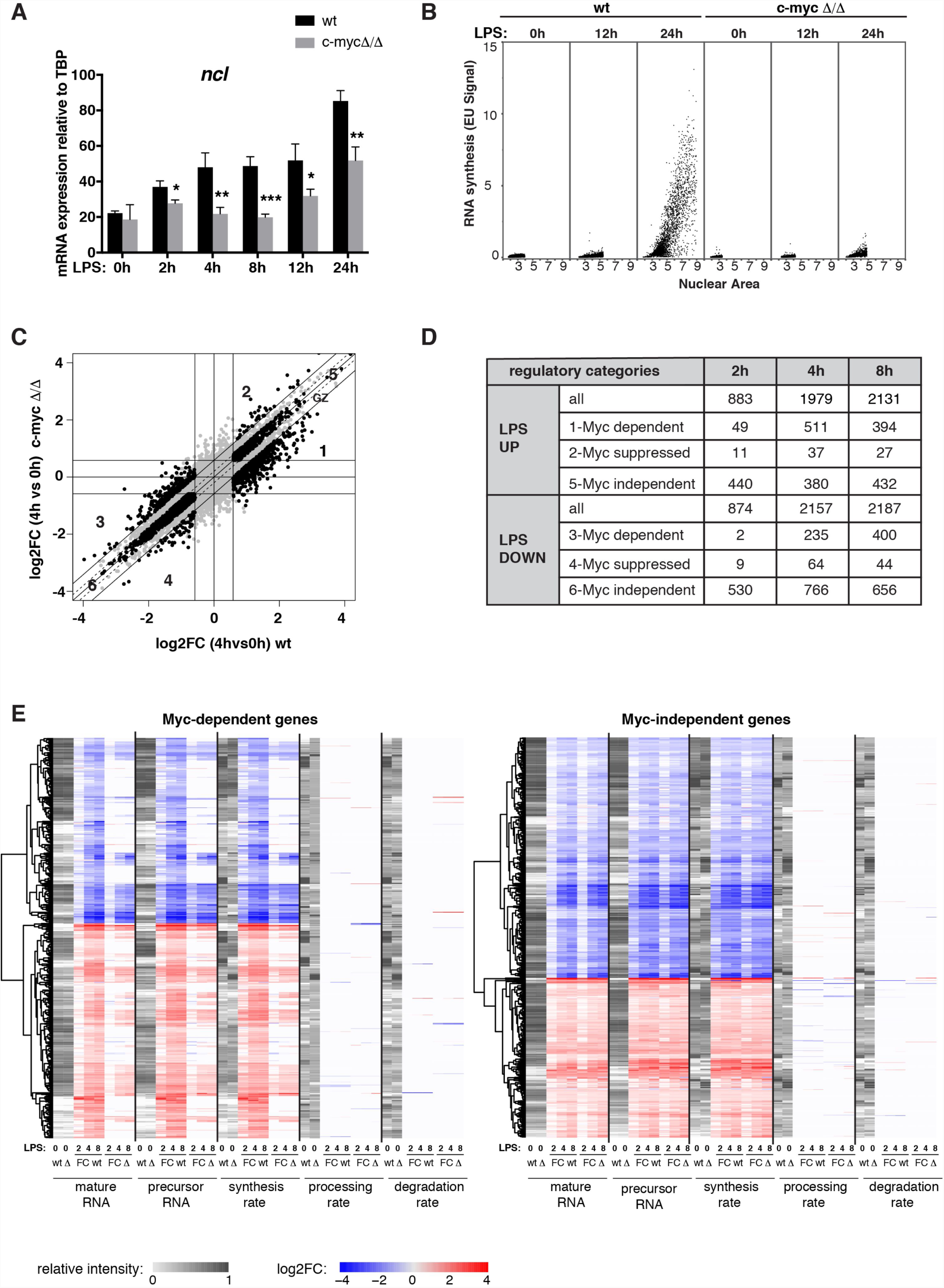
Myc is required for the regulation of a subset of LPS-responsive genes during B-cells activation. **A.** *Ncl* mRNA expression (normalized to *TBP*) at different time points after LPS stimulation in *c-myc*^*wt/wt*^ (in black) and c*-myc*^Δ/Δ^ cells (in gray). Data are presented as mean ± SD; n = 3. **B.** To quantify RNA synthesis at the single-cell level, we exposed our cultures to brief pulses of ethyl-uridine (EU) and measured its incorporation into RNA by light microscopy. The scatter plots show the nuclear Area (x axis) versus the EU signal (y axis) in *c-myc*^*wt/wt*^ and c*-myc*^Δ/Δ^ cells at different time points after LPS treatment, as indicated. One representative experiment out of 5 is shown. **C.** Scatter plot showing the log2 fold change (log2FC) of each expressed mRNA at 4h of LPS treatment relative to untreated cells, for *c-myc*^*wt/wt*^ (x-axis) and c*-myc*^Δ/Δ^ cells (y-axis). Regulatory groups 1-6 are as defined in the text. **D.** Numbers of genes classified in the different regulatory categories on the basis of the RNA-seq data at the different time points upon LPS stimulation. **E.** Heatmaps showing the variations in mature and precursor RNAs, synthesis, processing and degradation rates (log2FC) for Myc dependent (left) and independent genes (right), as defined at either 4 or 8h after LPS treatment; the grey scale represents the starting level for each parameter in unstimulated cells.

We used RNA-seq to profile the Myc-dependent transcriptional response shortly after LPS stimulation (2, 4 and 8h): based on a ≥1.5 fold-change to call differentially expressed genes (DEGs: Log_2_FC ≥0.58; qval ≤ 0.05), LPS induced the rapid up- and down-regulation of several thousand mRNAs (**Fig. EV2a)**. We defined Myc-regulated genes as those for which the magnitude of the LPS response was reduced by at least 1.5-fold in *c-myc*^Δ*/*Δ^ relative to *c-myc*^*wt/wt*^ cells (groups 1-4, **Fig. 1c, d, Supplementary Table 1, 2**): among these, the most abundant were Myc-dependent LPS-induced and repressed genes, both showing dampened responses in *c-myc*^Δ*/*Δ^ cells (groups 1 and 3), while much fewer mRNAs showed reinforced responses (groups 2 and 4). In line with the kinetics of Myc accumulation, Myc-dependent responses were rare at 2h but increased at later time-points (**Fig. 1d, Fig. EV2b**). Moreover, leaving aside an intervening “grey zone”, significant fractions of all mRNAs showed Myc-independent up- or down-regulation by LPS (altered ≤1.15 fold in *c-myc*^Δ*/*Δ^ relative to *c-myc*^*wt/wt*^ cells; groups 5, 6; **Fig. 1c, d**, **Supplementary Table 1, 2**). Besides analyzing mature mRNA species, we used intronic reads to quantify pre-mRNAs and to computationally model rates of RNA synthesis, processing and degradation along the time-course: as observed following MycER^T2^ activation in fibroblasts (de Pretis, Kress et al., 2017), the changes in pre-mRNA and mRNA levels elicited by either LPS or *c-myc* deletion correlated with variations in synthesis rate, with no significant alterations in either processing or degradation (**Fig. 1e**). In conclusion, Myc was rapidly induced by LPS and modulated the transcriptional response of select groups of genes in early G1 (4-8h), preceding general effects on biomass accumulation and cell size (Sabò et al., 2014).

We then used ChIP-seq to profile Myc along the genome: approximately 2000 binding sites were detected in resting wild-type B-cells, rising to ca. 22000 after LPS stimulation (either 2, 4 or 8h), with consistent overlaps along the time-course (**Fig. 2a**). Most Myc-binding sites were proximal (−2 to +1 kb) to an annotated transcription start site (TSS), albeit the fraction of distal sites increased upon stimulation (from 12% at 0h to 45% at 8h). Remarkably, the progression of Myc-binding profiles upon LPS-stimulation was virtually overlapping with that seen *in vivo* when comparing control *c-myc*^*wt/wt*^ (C), pre-tumoral Eµ-*myc* transgenic B-cells (P) and lymphomas (tumor: T) (**Fig. 2b, c**), consistent with parallel increases in Myc levels (Sabò et al., 2014). Further comparison with histone mark profiles in *c-myc*^*wt/wt*^ B-cells (Sabò et al., 2014) showed that Myc-binding sites pre-existed in an active state characterized by elevated H3K4me3 and H3K4me1 at proximal and distal sites, respectively, with H3K27ac at both (**Fig. 2b-e**), indicative of active promoters and enhancers. While few of these regulatory regions (<20% and <5%, respectively) were bound by Myc in naïve B-cells (0h or C), most were targeted upon either LPS stimulation or tumor development (**Fig. 2f)**: those bound in resting conditions corresponded to high affinity sites that were also the most efficiently bound upon Myc activation (**Fig. 2g**) and showed the highest enrichments for Myc-binding consensus motifs (**Fig. 2h)**. Altogether, in line with previous observations in either B-cells (Guccione, Martinato et al., 2006, Lin et al., 2012, Nie et al., 2012, Sabò et al., 2014) or others (Guo, Li et al., 2014, Kress et al., 2016, Kress et al., 2015, Lin et al., 2012, Lorenzin et al., 2016, Soufi, Donahue et al., 2012, Walz et al., 2014, Zeid et al., 2018), acute accumulation of Myc upon LPS treatment led to its widespread association with pre-existing promoters and enhancers: this phenomenon, also termed “invasion” (Lin et al., 2012, Nie et al., 2012, Sabò et al., 2014), most probably occurs through low affinity, non sequence-specific interactions with genomic DNA. Together with the findings that Myc deletion only impacts a subset of the genes regulated by either LPS (this work) or serum (Perna et al., 2012), we infer that chromatin association cannot be systematically equated with productive regulatory engagement of the transcription factor onto genomic DNA (de Pretis et al., 2017, Kress et al., 2015, Muhar et al., 2018).

**Fig. 2.**
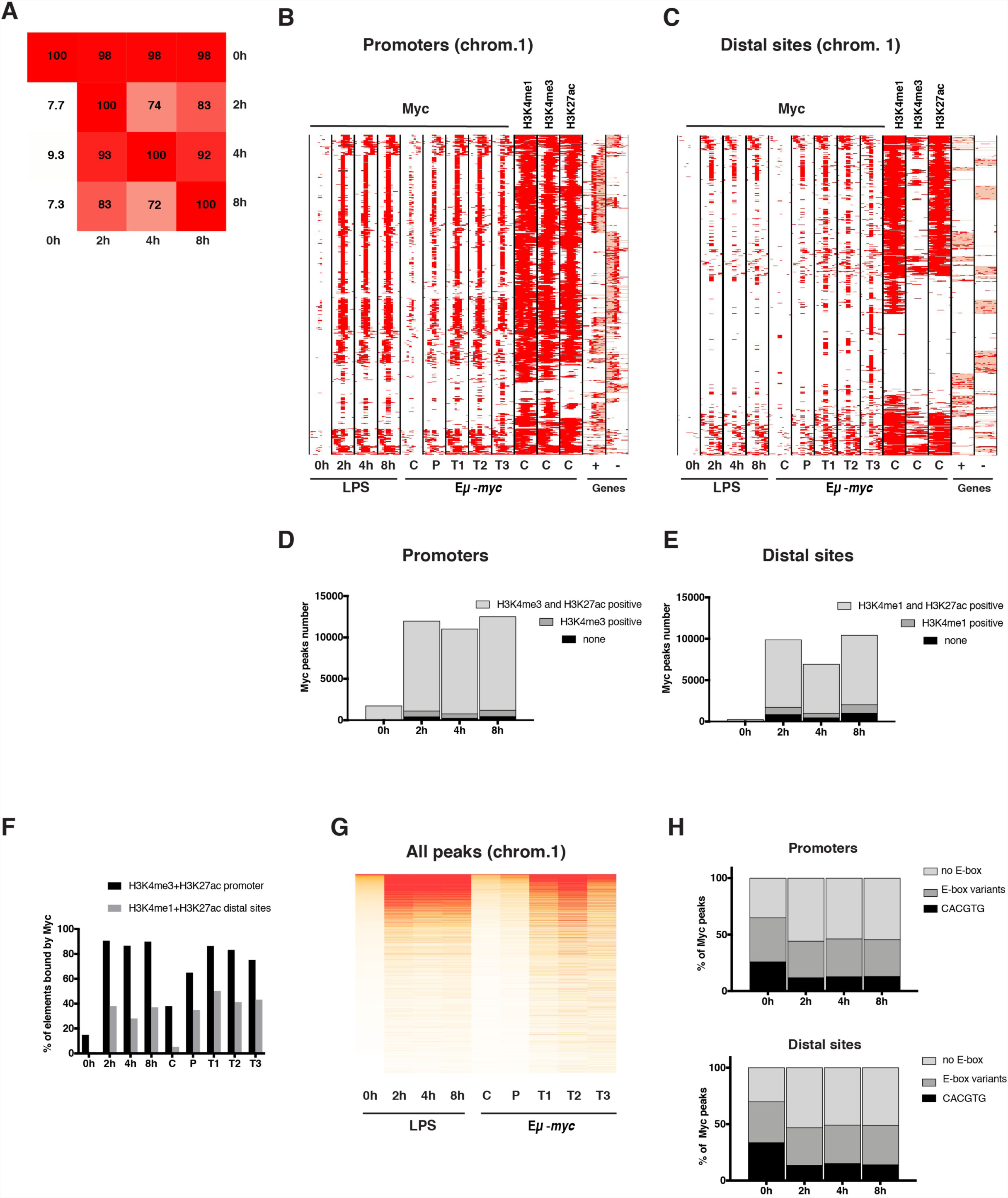
Myc widely associates with open chromatin upon LPS stimulation. **A.** Overlap between Myc ChIP-seq peaks. For each column, the percentage of peaks overlapping (over ≥1 bp) with the reference samples is reported. Peaks showed consistent distributions along the time-course, almost all those called in untreated cells being included in LPS-stimulated samples **B.** Heatmap showing the distribution of Myc at annotated promoters either *in vitro* in LPS-treasted B-cells (0, 2, 4 8h), or *in vivo* in control *c-myc*^*wt/wt*^ B-cells (C), pre-tumoral Eµ-*myc* B-cells (P) and tumors (T)(Sabò et al., 2014). Each row represents a different genomic interval (6 kb width, centered on Myc peaks). The panel includes every annotated promoter in chromosome 1 that was called as Myc-associated by ChIP-seq in at least one of the samples. For the same intervals, the distributions of H3K4me1, H3K4me3, and H3K27ac in the *in vivo* control sample, CpG Islands (CGIs) and annotated genes (exons in red, introns in pink; + sense, - antisense strand) are also shown. **C**. As in B., for distal Myc-binding sites. **D.** Bar plot representing the number of Myc peaks annotated at promoters (−2kb, +1kb) at different time points after LPS stimulation. The grey shadings mark the subsets of peaks associated with H3K4me3, with or without H3K27ac, as indicated. **E.** As in D, for distal Myc-binding sites, with H3K4me1 instead of H3K4me3. **F.** Bar plot representing the percentage of active promoters (positive for H3K4me3 and H3K27ac) or distal sites (H3K4me1 and H3K27ac) that overlap with at least a Myc peak at the different time points after LPS stimulation (0, 2, 4, 8h) or *in vivo* (C, P, T1, T2, T3), as indicated. **G.** Quantitative heatmap showing Myc signal intensities (read counts normalized by library size), ranked based on the control sample. **H.** Bar plot representing the percentage of Myc peaks at promoters (top) or distal sites (bottom) that contain canonical (CACGTG) or variant (CACGCG, CATGCG, CACGAG, CATGTG) E-boxes in a 200 bp window centered on the peak summit.

One parameter predicting transcriptional responses in other cell types was the relative efficiency in Myc binding at promoters (de Pretis et al., 2017, Kress et al., 2016, Lorenzin et al., 2016, Muhar et al., 2018), or “Myc share”(de Pretis et al., 2017): indeed, those genes for which the RNA synthesis rate increased upon LPS stimulation also showed the highest gains in Myc binding (i.e. increasing share), while those with a reduced synthesis showed the lowest gains (decreasing share) (**Fig. 3a, b**). This correlation was the strongest for Myc-dependent genes, although still present at Myc-independent loci, as also observed in a model of Myc-driven liver cancer (de Pretis et al., 2017, Kress et al., 2016): albeit paradoxical at first sight, this observation is consistent with the notion that promoter activity and Myc binding are mutually dependent, owing to the interaction of Myc with open chromatin, co-factors and components of the basal transcription machinery (Guccione et al., 2006, Richart, Carrillo-de Santa Pau et al., 2016, Thomas, Wang et al., 2015). In previous work, high-affinity E-box-containing loci were deemed to be already saturated by Myc at the population level in proliferating U2OS cells, thus showing no – or negligible – increases in either Myc binding or transcriptional activity upon Myc overexpression (Lorenzin et al., 2016): in our experiments instead, virtually all promoters – regardless of initial binding intensities – showed increased Myc binding upon stimulation (**Fig. 3c**), implying that these were far from saturation to start with. Accordingly, Myc-dependent LPS-induced genes, showed not only the highest increase in Myc binding (**Fig. 3a**) but also the highest frequency of E-boxes (**Fig. 3d**). Hence, upon B-cell stimulation, with a concomitant transition from very low to high Myc levels, Myc drove rapid and selective activation of high-affinity promoters, most frequently – but not always – associated with the presence of the E-box binding motif.

**Fig. 3.**
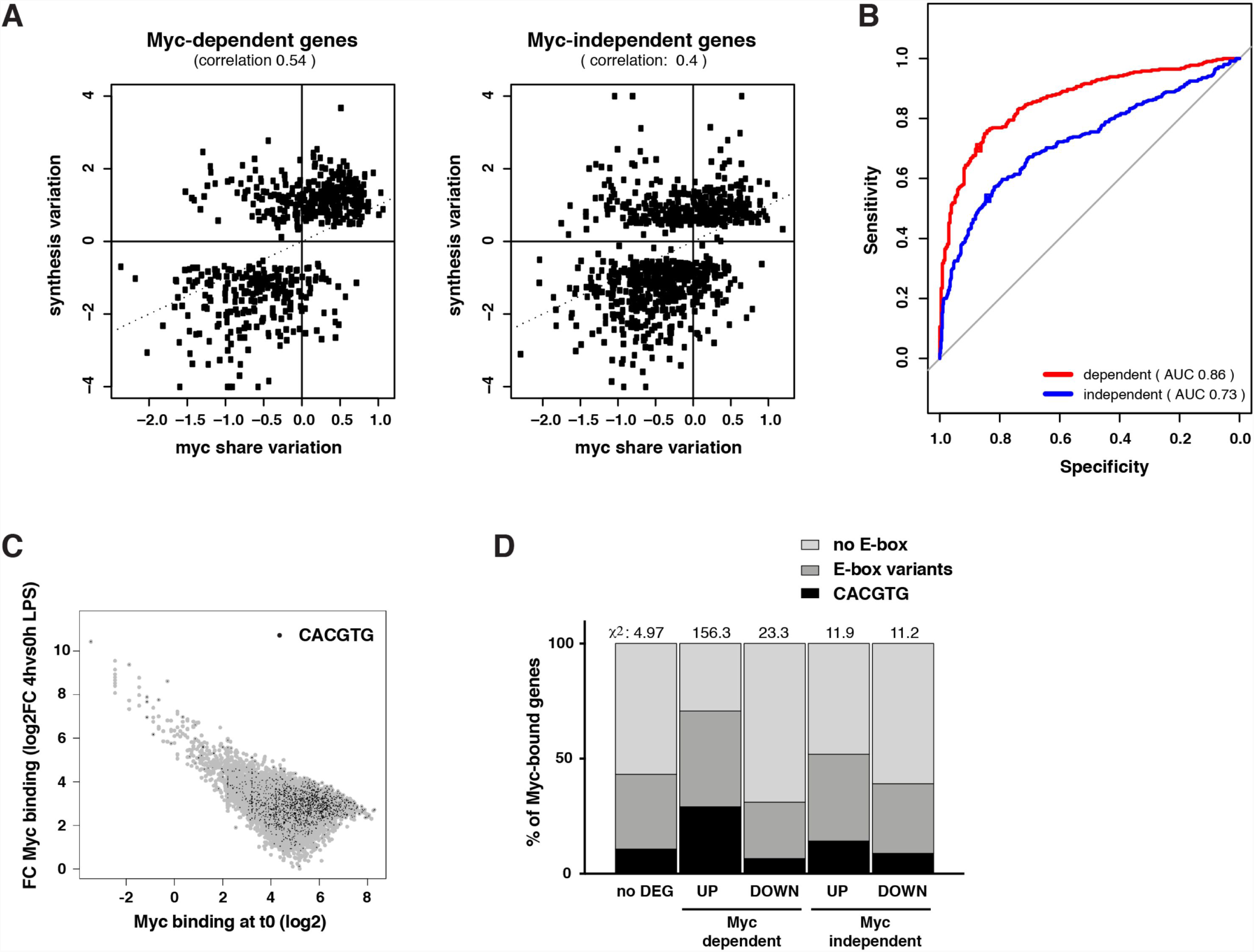
Changes in Myc share are predictive of gene regulation. **A.** Scatter plots correlating the variations in Myc share at promoters (averaged between the 2, 4 and 8h time points) and in RNA synthesis (at 8h), relative to untreated cells, for Myc-dependent (left) and independent genes (right). The Spearman correlation between the two parameters is reported. **B.** Receiver operating characteristic (ROC) curves for the ability of discriminating induced and repressed genes at increasing thresholds of changes in Myc binding. AUC = Area under the curve. For each system, the dot corresponds to the variation of Myc at which promoters begin increasing their share of Myc binding. **C.** Scatter plot representing the fold-change in Myc binding at 4h LPS (relative to time zero) as a function of the initial binding intensity at time zero (expressed as log2 of the coverage in a 200 bp window around the summit of the peak) for each promoter bound by Myc at 4h. Peaks containing a canonical E-box are highlighted in black. **D.** Bar plot showing the percentage of Myc-bound promoters within the indicated transcriptionally regulatory categories that contains a canonical or variant E-box. The overall chi-square is 207.7 with a p-value <0.00001. The contribution to the chi-square of each category is reported above the corresponding bar in the plot.

In order to address the mechanisms of transcriptional regulation by Myc, we profiled RNA polymerase II (RNAPII) by ChIP-seq, computed its quantitative changes in the promoter (TSS), gene body (GB) and termination site (TES) of regulated genes, and confronted those to the changes in RNA synthesis rates (**Fig. EV3a, b**). As expected, LPS-induced genes showed consistent increases in RNAPII densities in all of their domains, the opposite being true for repressed genes: most importantly, these effects of LPS on RNAPII were suppressed by *c-myc* deletion at Myc-dependent, but not Myc-independent loci (**Fig. EV3 c, d**). We then used a dedicated algorithm (de Pretis et al., 2017) to model the kinetic rates governing the RNAPII cycle at each locus, including its recruitment to the promoter (p_1_), pause-release (p_2_), elongation (p_3_) and release from the transcription end site (TES, p_4_) (**Fig. 4a**). Remarkably, the four rates were altered at Myc-dependent genes in *c-myc*^Δ*/*Δ^ cells, indicating that Myc deletion impacted on all steps of the transcription cycle (**Fig. 4b, c**): however, as seen after MycER^T2^ activation in fibroblasts(de Pretis et al., 2017), RNAPII recruitment was the most significantly affected step upon either LPS stimulation or Myc deletion. To increase the resolution of our analysis, we clustered LPS-regulated genes on the basis of RNAPII dynamics (**Fig. 4d, e;** and **Fig. EV4, 5, Supplementary Table 3**): remarkably, the largest sets of LPS-activated genes (CL1: Myc-dependent; CL9: Myc-independent), were almost exclusively regulated through RNAPII loading (**Fig. 4f, h**) while others showed significant contributions from other regulatory steps (**Fig. EV4, 5**). Among the Myc-dependent induced genes, those of CL1 also showed the highest increases in Myc share (**Fig. 4g**). Myc-dependent repressed genes, instead (CL3, 6, 7, 8), were characterized by a relative loss of Myc binding (decreased share) and by decreased RNAPII recruitment (**Fig. 4b, Fig. EV4**), suggesting that Myc-dependent repression could largely be a passive process(de Pretis et al., 2017) – albeit not excluding the existence of active repressive mechanisms at select loci (Tu, Shiah et al., 2018, Walz et al., 2014). Altogether, while modulating all stages in the RNAPII cycle, Myc primarily drives RNAPII recruitment at activated loci (de Pretis et al., 2017): while it was suggested that Myc mainly regulates pause-release (Rahl, Lin et al., 2010), our work and others (Davari, Lichti et al., 2017, de Pretis et al., 2017) indicate that a careful integrative analysis of RNAPII and RNA dynamics is needed to unravel the hierarchical contribution of distinct regulatory steps.

**Fig. 4.**
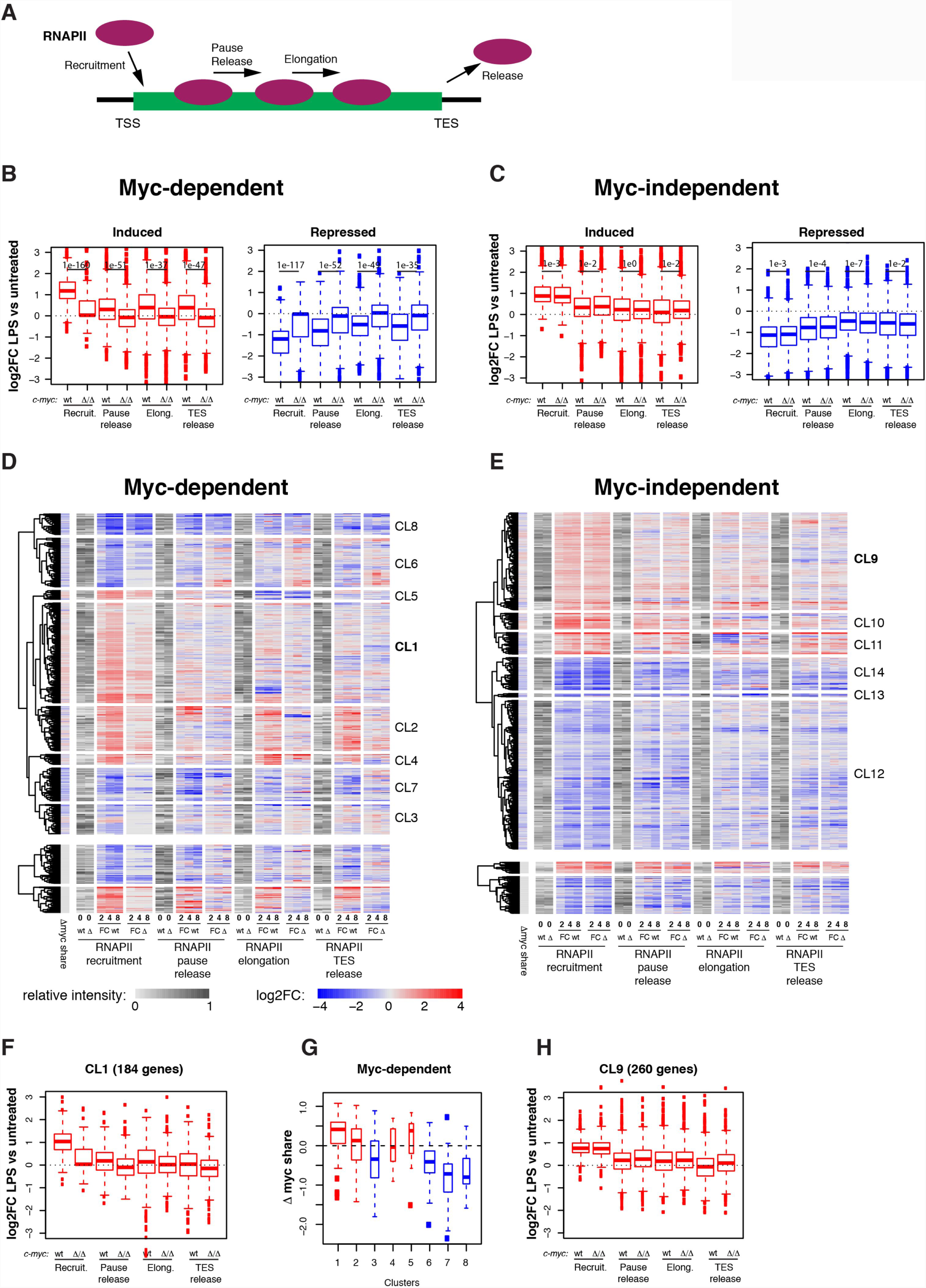
Myc regulates all steps of the RNAPII transcriptional cycle. **A.** Scheme depicting the different steps of RNAPII regulation for which the corresponding rates were modelled. Changes in density of RNAPII in each region of the transcriptional unit are considered as the net result of the RNAPII coming from the upstream region, and that exiting toward the downstream region, both governed by the corresponding kinetic rate (de Pretis et al., 2017). **B.** Boxplots reporting LPS-induced changes (Log2FC) for each of the four RNAPII kinetic rates in *c-myc*^*wt/wt*^ (wt) and c*-myc*^Δ/Δ^ cells for Myc-dependent induced and repressed genes, as indicated. p-values obtained with two-sample Wilcoxon tests for the comparison between *c-myc*^*wt/wt*^ and c*-myc*^Δ/Δ^ cells are reported **C.** As in B, for Myc-independent genes. **D.** Heatmap showing the variations in RNAPII kinetic rates (log2FC) on Myc-dependent genes during the LPS time course. The grey scale represents the starting level for each parameter in unstimulated cells. The first column on the left reports the changes in Myc share for the same genes. Genes were clustered using hierarchical clustering. **E.** As in D, for Myc-independent genes. **F.** As in B, for the genes of cluster 1. **G.** Boxplots reporting the changes in Myc share in LPS treated versus untreated cells for the different clusters of panel d. **H.** As in C, for the genes of cluster 9.

To discern the biological pathways directly modulated by Myc, we analyzed the gene ontology (GO) terms enriched in the various subgroups of LPS-induced genes: Myc-dependent (or Group 1, **Fig. 1c, Supplementary Table 2**: 647 genes), Myc-independent (Group 5: 697 genes) and the intervening “grey zone” (GZ: 1129 genes) as the latter was likely to comprise genes modulated by Myc with low – albeit possibly biologically meaningful – amplitude. We also analysed Cluster 1, characterized by the highest increase in Myc share (CL1, **Fig. 4d**, 184 genes). Group 1, GZ and CL1 most significantly enriched for common GO terms pertaining to RNA and amino acid metabolism, as well as mitochondrial biogenesis (**Supplementary Table 4**), with additional terms enriched only in the GZ, such as translation, ribosome biogenesis, RNAPI transcription, or nucleotide biosynthesis. Importantly, Myc-independent genes showed little overlap with any of the above, enriching for distinct functional categories such as lymphocyte activation, cell cycle, apoptosis, DNA metabolism (repair, recombination). Thus, biological processes that depend upon Myc activity (e.g. cell cycle, DNA replication) are regulated – to a large extent – by genes that show Myc-independent expression, suggesting that Myc acts on essential upstream events (e.g. protein and nucleic acid biosynthesis). Most importantly in this context, other transcription factors such as E2F or NFY may have predominant roles – or be redundant with Myc (Liu, Tang et al., 2015) – in regulating Myc-independent genes (**Fig. EV6**).

It is noteworthy here that several of the Myc-dependent ontological categories (e.g. RNA metabolism, ribosome biogenesis, nucleotide biosynthesis, etc…) were also enriched among Myc-regulated genes in different cell types (Lorenzin et al., 2016, Muhar et al., 2018, Perna et al., 2012), in germinal center B-cells (Dominguez-Sola et al., 2012), or in primary B-cells at longer time points of LPS stimulation (72h)(Perez-Olivares et al., 2018). Yet, the overlaps between these and our gene lists were only modest (**Fig. EV7a, Supplementary Table 5**). Hence, while different genes may show Myc-dependent expression in different contexts, the biological processes that rely on Myc activity appear to be generally conserved. Remarkably, larger fractions of the Myc-dependent genes identified in this work were induced during lymphomagenesis in Eµ-*myc* transgenic mice (Sabò et al., 2014) (**Fig. EV7b**), emphasizing the need to confront normal and pathological Myc-regulated programs in the corresponding cellular contexts.

Altogether, we have identified approximately 650 genes that were induced in a Myc-dependent manner within 4-8h following stimulation, and an additional group of ca. 1100 additional genes regulated by Myc with modest quantitative effects: while the latter were below threshold for being reliably called as Myc-dependent, both groups enriched for highly consistent functional categories. By and large, Myc-dependent LPS-responsive genes encode proteins involved in RNA biology, energy production and anabolic pathways (Caro-Maldonado et al., 2014, Dang, 2013, Wang et al., 2011): these, in turn, may provide the building blocks (nucleotides, amino acids) and energy required to sustain the large accumulation of biomass – and in particular RNA – characteristic of activated B-cells (Kieffer-Kwon et al., 2017, Nie et al., 2012, Sabò et al., 2014). In an alternative model, Myc was deemed to act as a direct activator – or *amplifier* – of all expressed genes (Lin et al., 2012, Nie et al., 2012, Porter et al., 2017, Zeid et al., 2018). However, our results and others (Muhar et al., 2018, Perna et al., 2012, Sabò et al., 2014, Walz et al., 2014) provide no formal support for this model: instead, when observed, RNA amplification is most consistently interpretable as a late, indirect consequence of Myc action, mediated by a selective – yet complex – set of target genes (Kress et al., 2015, Sabò & Amati, 2018). Most importantly, the Myc-dependent program identified here in activated B-cells was constitutively deregulated during lymphomagenesis in Eµ-*myc* transgenic mice (Sabò et al., 2014). Hence, besides aberrant regulation of novel, tumor-specific targets (Lorenzin et al., 2016, Sabò et al., 2014, Walz et al., 2014), the oncogenic action of Myc may largely rely on the uncontrolled activation of genes regulated during mitogenic stimulation in normal cells.

## Materials & Methods

### Mouse strains and primary B cells

C57BL/6 *c-myc*^*f/f*^ mice were obtained from Andreas Trumpp (Trumpp et al., 2001). Naïve mouse B-cells were isolated from the spleen of 7–10 weeks old wild type or *c-myc*^*f/f*^ mice with the B-cell isolation kit (MACS Miltenyi Biotec Cat. no. 130-090-862). After purification, naïve B cells were incubated for 1h at 37°C with recombinant Tat-Cre protein (50 μg/ml) in optimem + 1% fetal bovine serum (Peitz, Pfannkuche et al., 2002) in order to induce deletion of the c-*myc*^*f/f*^ allele. Cells were washed with PBS and then grown in B cell medium composed of DMEM medium (Dulbecco’s Modified Eagle Medium) and IMDM medium (Iscove’s Modified Dulbecco’s Medium) in a 1:1 ratio and additionated of 10% fetal calf serum (FCS) (Globefarm Ltd, Cranleigh, UK), 2 mM L-glutamine (Invitrogen Life Technologies, Paisley, UK), 1% non-essential amino acids (NEAA), 1% penicillin/streptomycin and 25 μM β-mercaptoethanol (Gerondakis et al., 2007). 12h after seeding, B cells were stimulated with lipopolysaccharide LPS (50 μg/ml; SIGMA L6237) to induce cell activation. The Tat-Cre protein was produced and purified as previously described (Peitz et al., 2002).

### Proliferation, cell cycle analysis and sorting

For proliferation analysis, cells were counted with Trypan Blue to exclude dead cells every 24h. For cell cycle analysis, cells were incubated with 33 μM BrdU for a pulse labelling of 30 min. Cells were then harvested, washed with PBS and fixed in ice-cold ethanol. Upon DNA denaturation using 2N HCl, cells were stained with an anti-BrdU primary antibody (BD Biosciences) and anti-mouse FITC conjugated secondary antibody (Jackson Immunoresearch). DNA was stained by resuspending the cells in 2.5 μg/ml Propidium Iodide (Sigma) overnight at 4°C before FACS analysis. All samples were acquired on a FACS Canto II (BD Biosciences) flow cytometer. At least 15,000 events were acquired and the analysis was performed using the FlowJo X software. For cell sorting, cells were resuspended in cold Macs Buffer (0.5 % BSA, 2 mM EDTA in PBS) and sorted on the basis of the FSC/SSC parameters with a Facs-Aria II machine (BD Biosciences).

### Immunoblot analysis

5×10^6^ B-cells were lysed with RIPA Buffer (20 mM HEPES at pH 7.5, 300 mM NaCl, 5 mM EDTA, 10% Glycerol, 1% Triton X-100, supplemented with protease inhibitors (Roche) and phosphatase inhibitors (0.4 mM Orthovanadate, 10 mM NaF) and briefly sonicated. Cleared lysates were electrophoresed and immunoblotted with the indicated primary antibodies: c-Myc Y69 (ab32072) from Abcam, Vinculin (V9264) from Sigma. Chemioluminescent detection, after incubation of the membranes with appropriate secondary antibodies, was done through a CCD camera using the ChemiDoc System (Bio-Rad). Quantification of protein levels was done using the Image Lab Software (Bio-Rad, version 4.0).

### EU staining and data analysis

In order to label newly synthesized RNA, B-cells were plated at a cell density of 8*10^5/ml and incubated with the alkyne-modified nucleoside, 5-ethynyl uridine (EU), 1 mM for 1h before fixation in 4% PFA for 10 min at RT. Fixed cells were washed in PBS and resuspended in PBS+BSA 3%. Cells were then cytospinned on polylysine-coated slides, permeabilized with Triton X-100 0.5% in PBS and treated with the Click-iT reaction cocktail for 30 min at RT as indicated by manufacturer’s instruction (Click-iT RNA Imaging Kit – Invitrogen, C10329). DNA was than stained with DAPI.

The image analysis was performed using a custom pipeline developed and executed in the Acapella software development/run-time environment (Perkin Elmer). Nuclei where detected on the basis of DAPI staining using a Perkin Elmer proprietary algorithm and each nucleus was associated to a nuclear area and an integrated EU signal.

### Isolation of genomic DNA

Cells pellet (1.5*10^6) were collected at different time points after LPS stimulation and DNA was extracted with the Nucleospin tissue kit (Macherey-Nagel, 740952). The genomic DNA was eluted in 50 μl of BE buffer (5 mM Tris/HCl, pH 8.5). The analysis of *c-myc* deletion efficiency was performed by real-time PCR using 10 ng of genomic DNA as template and the primers reported in **Supplementary Table 6** (Trumpp et al., 2001).

### RNA extraction and analysis

Total RNA (at least from 2.5* 10^6 cells) was purified onto RNeasy columns (Qiagen) and treated on-column with DNase (Qiagen). Complementary DNA (cDNA) was prepared using ImProm-II™ reverse transcription kit (Promega, A3800) and 10 ng of cDNA were used as template for each real-time PCR reaction. cDNA was detected by fast SyberGreen Master Mix (Applied Biosystems, 4385614) on CFX96 Touch™ Real-Time PCR Detection System (Biorad). Sequences of the used PCR primers were reported in **Supplementary Table 6**.

For RNA-seq experiments, total RNA from 8^10^6^ B-cells was purified as above, then 0.5 μg were treated with Ribozero rRNA removal kit (Epicentre) and EtOH precipitated. RNA quality and removal of rRNA were checked with the Agilent 2100 Bioanalyser (Agilent Technologies). Libraries for RNA-Seq were then prepared with the TruSeq RNA Sample Prep Kits v2 (Illumina) following manufacturer instruction (except for skipping the first step of mRNA purification with poly-T oligo-attached magnetic beads). RNA-seq libraries were then run on the Agilent 2100 Bioanalyser (Agilent Technologies) for quantification and quality control and then sequenced on Illumina HiSeq2000.

### Chromatin Immunoprecipitation

Purified splenic B-cells were resuspended in PBS at room temperature and fixed for 10 min by addition of formaldehyde to a final concentration of 1%. Fixation was stopped by addition of glycine to a final concentration of 0.125 M. Cells were washed in PBS, resuspended in SDS buffer (50 mM Tris at pH 8.1, 0.5% SDS, 100 mM NaCl, 5 mM EDTA, and protease inhibitors) and stored at −80°C before further processing for ChIP as described in (Sabò et al., 2014). For ChIP-Seq analysis, lysates obtained from 30-50×10^6^ B-cells were immunoprecipitated with 10 μg of Myc (Santa Cruz, sc-764) or RNAPII (Santa Cruz, sc-899) antibodies. Immunoprecipitated DNA was eluted in TE-2% SDS and crosslinks were reversed by incubation overnight at 65 °C. DNA was then purified by Qiaquick columns (Qiagen) and quantified using Qubit™ dsDNA HS Assay kits (Invitrogen). 1.5-2 ng of ChIP DNA was end-repaired, A-tailed, ligated to the sequencing adapters and amplified by 17-cycles of PCR, size selected (200-300bp) according with TruSeq ChIP Sample Prep Kit (Illumina). ChIP-seq libraries were then run on the Agilent 2100 Bioanalyser (Agilent Technologies) for quantification and quality control and then sequenced on the Illumina HiSeq2000. Sequences of the primers used in qPCR were reported in **Supplementary Table 6**.

### Next generation sequencing data filtering and quality assessment

ChIP-seq and RNA-seq reads were filtered using the fastq_quality_trimmer and fastq_masker tools of the FASTX-Toolkit suite (http://hannonlab.cshl.edu/fastx_toolkit/). Their quality was evaluated and confirmed using the FastQC application: (www.bioinformatics.babraham.ac.uk/projects/fastqc/). Pipelines for primary analysis (filtering and alignment to the reference genome of the raw reads) and secondary analysis (expression quantification, differential gene expression and peak calling) have been integrated in the HTS-flow system (Bianchi, Ceol et al., 2016). Bioinformatic and statistical analysis were performed using R with Bioconductor and comEpiTools packages (Gentleman, Carey et al., 2004, Kishore, de Pretis et al., 2015).

### ChIP-seq data analysis

ChIP-Seq NGS reads were aligned to the mm9 genome through the BWA aligner (Li & Durbin, 2009) using default settings. Peaks were called using the MACS software (v2.0.9)(Zhang, Liu et al., 2008) with the option ‘– mfold = 7,30 -p 0.00001’, thus outputting only enriched regions with P-value <10-5. Normalized read counts within a genomic region were determined as the number of reads per million of library reads (total number of aligned reads in the sequencing library). Peak enrichment was determined as log2(Peakw/Nc — inputw/Ni), where Peakw is the read count on the enriched region in the ChIP sample, inputw the read count on the same region in the corresponding input sample, Nc is the total number of aligned reads in the ChIP sample, and Ni is the total number of aligned reads in the input sample. Promoter peaks were defined as all peaks with at least one base pair overlapping with the interval between − 2 kb to +1 kb from the nearest TSS. For Myc share calculation promoters were defined as − 2 kb to +2 kb from the nearest TSS. For RNAPII ChIP-seq analysis the following genomic regions were considered: promoters (−50, +700 from TSS); TES (−1kb,+4kb from TES), gene body (from promoter end to TES start). The presence of canonical and variant E-boxes (CACGCG, CATGCG, CACGAG, CATGTG)(Blackwell, Kretzner et al., 1990, Grandori, Mac et al., 1996, Perna et al., 2012) in the Myc ChIP peaks was scored in a region of 200 bp around the peak summit. The Myc share at each promoter was determined by normalizing the ChIP-seq signal by the total amount of promoter-bound Myc in the genome at the same time-point.

When comparing Myc ChIP-seq with H3K4me3, H3K4me1 or H3K27ac histone marks to define peaks in active promoter or enhancers (Calo & Wysocka, 2013, Zhou, Goren et al., 2011), we considered two peaks as overlapping when sharing at least one base pair (subsetByOverlaps tool of the comEpiTools R package).

Fold change of Myc binding at promoters was calculated as the log2 ratio of the coverage in a 200 bp window around the summit of the peaks identified at 4h of LPS in stimulated versus not stimulated conditions. To avoid regions with 0 coverage in the unstimulated situation, all those regions were set to the minimum coverage identified.

### RNA-seq data analysis

RNA-Seq NGS reads were aligned to the mm9 mouse reference genome using the TopHat aligner (version 2.0.8)(Kim, Pertea et al., 2013) with default parameters. In case of duplicated reads, only one read was kept. Read counts were associated to each gene (based on UCSC-derived mm9 GTF gene annotations), using the featureCounts software (http://bioinf.wehi.edu.au/featureCounts/) (Liao, Smyth et al., 2014) setting the options -T 2 -p -P. Absolute gene expression was defined determining reads per kilobase per million mapped reads defining total library size as the number of reads mapping to exons only (eRPKM). After removing very low expressed genes (below 1 eRPKM in all samples), we obtained a set of 12690 expressed genes that were used for further analysis. Differentially expressed genes (DEGs) were identified using the Bioconductor Deseq2 package (Love, Huber et al., 2014) as genes whose q-value is lower than 0.05 and |FoldChange|>1.5. The different categories of Myc-dependent LPS response and Myc-independent LPS response, were defined at each time point (t_i_) relative to time zero (t0), as follows:

p_x_=qval t_i_ vs t0 (*c-myc*^*wt/wt*^)

p_y_=qval t_i_ vs t0 (c*-myc*^Δ/Δ^)

x=log2FC t_i_ vs t0 (*c-myc*^*wt/wt*^)

y=log2FC t_i_ vs t0 (c*-myc*^Δ/Δ^)

- <u>NO DEG</u>: (p_x_ > 0.05 & p_y_ > 0.05) OR ((|x|≤ log2(1.5) & (|y|≤ log2(1.5)))
- <u>Class 5 (Myc-independent UP)</u>: (p_x_ ≤ 0.05 & p_y_ ≤ 0.05) & (x > log2(1.5) & y > log2(1.5)) & (|y-x|≤ log2(1.15))
- <u>Class 6 (Myc-independent DOWN)</u>: (p_x_ ≤ 0.05 & p_y_ ≤ 0.05) & (x < -log2(1.5) & y < -log2(1.5)) & (|y-x|≤ log2(1.15))
- <u>Class 1 (Myc-dependent UP)</u>: (p_x_ ≤ 0.05) & (x > log2(1.5)) & (y-x<−log2(1.5))
- <u>Class 3 (Myc-dependent DOWN)</u>: (p_x_ ≤ 0.05) & (x < -log2(1.5)) & (y-x>log2(1.5))
- <u>Class 2 (Myc-suppressed UP)</u>: (p_x_ ≤ 0.05 & p_y_ ≤ 0.05) & (x > log2(1.5)) & (y-x>log2(1.5))
- <u>Class 4 (Myc-suppressed DOWN)</u>: (p_x_ ≤ 0.05 & p_y_ ≤ 0.05) & (x < -log2(1.5)) & (y-x<−log2(1.5))

Functional annotation analysis to determine enriched Gene Ontology was performed using the ClueGo v2.5.3 Application of CytoScape v3.7.1 with the following parameters:

- Statistical Test Used = Enrichment/Depletion (Two-sided hypergeometric test)
- Correction Method Used = Bonferroni step down
- Min GO Level = 6
- Max GO Level = 8
- GO Fusion = true
- Show only pathways with p-value≤ 0.05
- GO Group = true
- Kappa Score Threshold = 0.4
- Group By Kappa Statistics = true
- Initial Group Size = 1
- Sharing Group Percentage = 50.0
- Merge redundant groups with >50.0% overlap

GO categories enriched in at least one gene list were reported in **Supplementary Table 4** and manually grouped on the basis of the ClueGo assignation to different groups.

Analysis of cis-regulatory motif enrichment in the promoter of the different classes of genes has been performed with the online tool available at http://software.broadinstitute.org/gsea/msigdb/index.jsp.

For the calculation of a significant overlap between Myc-dependent LPS-induced genes and the datasets of interest a hypergeometric distribution was assumed.

The external data were converted in the mouse orthologous when required and filtered keeping only the genes expressed in the reference dataset (B-cells, this study). Gene lists are reported in **Supplementary Table 5** and obtained from the following sources:

> Column D: MYC/MAX KO B-cells (Perez-Olivares et al., 2018)
>
> Column E: GFP-MYC+ GC B-cells (Dominguez-Sola et al., 2012)
>
> Column F: Myc-dependent serum responsive (Perna et al., 2012)
>
> Column G: Myc-dependent in U2OS cells (Lorenzin et al., 2016) (qval<0.01; log2FC<−1)
>
> Column H: Myc-dependent in K562 cells (Muhar et al., 2018) (qval<0.05; log2FC<−1)
>
> Column I: Myc-dependent in HCT116 cells (Muhar et al., 2018) (qval<0.05; log2FC<−0.5)
>
> Column J: Induced in Pre-tumoral Eµ-*myc* B-cells (Sabò et al., 2014) (qval<0.05; log2FC>0.585)
>
> Column K: Induced in Eµ-*myc* tumors (Sabò et al., 2014) (qval<0.05; log2FC>0.585)

### Estimation of synthesis, processing and degradation rates

Our previous R/Bioconductor package INSPEcT (de Pretis, Kress et al., 2015) was designed to estimate the rates of RNA synthesis, processing and degradation following metabolic labeling and quantification of total and newly synthesized RNA: we recently extended this package to pursue the same goal in the absence of experimental data on newly synthesized RNA, taking advantage of intronic reads in total RNA-seq experiments (INSPEcT2: Furlan et al., in preparation). Myc-dependent and independent genes at either 4 of 8h following LPS stimulation (**Supplementary Table 2**) were used for the modeling after removal of genes without introns or with scarce intronic signal. Out of the 925 Myc-dependent genes modeled, 674 were identified as modulated in their synthesis rate in the *c-myc*^*wt/wt*^ condition and kept for subsequent analysis. Similarly, 1005 Myc-independent genes out of the 1660 modeled were identified as modulated in their synthesis rates both in the *c-myc*^*wt/wt*^ and *c-myc*^Δ*/*Δ^ conditions and kept for subsequent analysis.

### Estimation of RNAPII recruitment, pause-release, elongation and termination rates

Similarly to what done in previous work(de Pretis et al., 2017), we modeled RNAPII progression on each gene as a dynamic system composed by 4 steps: recruitment (*p*_1_), pause-release (*p*_2_), elongation (*p*_3_), and the release from termination-end-sites (release, *p*_4_). This system relates to RNAPII quantification at promoters (*R*_*tss*_), gene-body (*R*_*gb*_) and termination end sites (*R*_*tes*_) as follows:

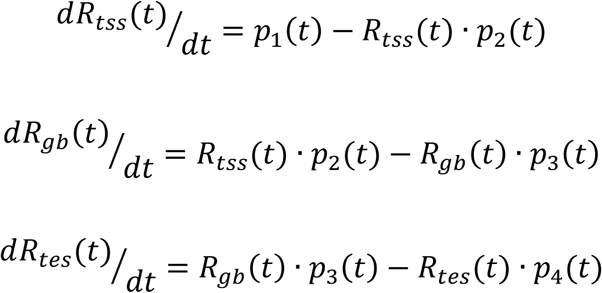

In practical terms, this model assumes that:

- RNAPII at transcriptional start site (*R*_*tss*_) is the balance between the amount of recruited RNAPII (*p*_1_(*t*)) and the amount of RNAPII that enters the gene-body due to pause-release (*R*_*tss*_(*t*) · *p*_2_(*t*));
- RNAPII at the gene-body (*R*_*gb*_) is the balance between the amount of RNAPII that enters the gene-body due to pause-release and the amount that enters into the termination-sites due to elongation (*R*_*gb*_(*t*) · *p*_3_(*t*));
- RNAPII at the termination-sites (*R*_*tes*_) is the balance between the amount of RNAPII that enters into the termination sites due to elongation and the amount that is released to nucleoplasm (*R*_*tes*_(*t*) · *p*_4_(*t*)).

Additionally, we assumed that the RNAPII that enters in termination-sites after elongation has synthesized an RNA molecule:

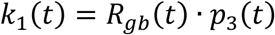

Estimated values of *p*_1_(*t*), *p*_2_(*t*), *p*_3_(*t*) and *p*_4_(*t*) are obtained at each experimental time point (0h, 2h, 4h, 8h) directly by the solution of the system of the four equations above, where *dR*_*tss*_(*t*)/*dt, dR*_*gb*_(*t*)/*dt*, and *dR*_*tes*_(*t*)/*dt* are estimated from the time course of *R*_*tss*_, *R*_*gb*_ and *R*_*tes*_, respectively, and *k*_1_(*t*) is the synthesis rate calculated by INSPEcT at the time *t*.

The relative error of each modeled gene is calculated as:

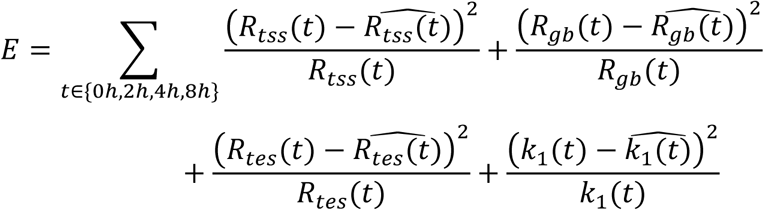

where 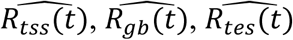, and 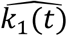 are the modeled values of *R*_*t*s*s*_(*t*), *R*_*gb*_(*t*), *R*_*tes*_(*t*), and *k*_1_(*t*) obtained by integrating the estimated rates *p*_1_(*t*), *p*_2_(*t*), *p*_3_(*t*) and *p*_4_(*t*) into the system of differential equations, assuming linear behavior of the estimated rates between the experimental time points. We considered valid only those models with relative error lower than 1 and no missing-values in any of the rates at any time point (618/674 Myc-dependent genes and 891/1005 Myc-independent genes).

## Statistical analysis

All the experiments, except for ChIP, were performed at least in biological triplicates. Two tailed-Student t-test was used to compare between two groups and expressed as p-values. In the figures: *P < 0.05, **P < 0.01, ***P < 0.001.

### Code availability

All R scripts used in data analysis and generation of figures are available upon request.

### Data availability

RNA-seq and ChIP-seq data have been deposited in NCBI’s Gene Expression Omnibus (GEO) and are accessible through GEO series accession number GSE126340.

### Materials & Correspondence

Correspondence and requests for materials should be addressed to A.S. or B.A.

## Acknowledgements

We thank P. Nicoli, A. Gobbi and M. Capillo for their help with the management of mouse colonies, S. Bianchi, L. Rotta and T. Capra for assistance with the Illumina HiSeq, S. Barozzi and D. Parazzoli for assistance with imaging technologies, A. Bisso, S. Campaner, O. Croci, G. Natoli and J. Zuber for discussions, comments and advice on data analysis. A.S. and B.A. acknowledge the support from the Istituto Italiano di Tecnologia in the early phases of the work. This study was supported by funding from the European Research Council (grant agreement no. 268671-MYCNEXT), the Italian Health Ministry (RF-2011-02346976) and the Italian Association for Cancer Research (2012-13182 and 2015-16768) to B.A., and from Worldwide Cancer Research (15-1260) to A.S.

## Author contributions

A.S. and B.A. designed experiments and wrote the manuscript. A.T. performed the experiments. A.V. and M.D provided technical support. A.A. analyzed the EU staining. S.d.P., M.F., M.F., M.J.M., M.P. and A.S. were involved in bioinformatic data analysis.

## Competing interests

The authors declare that they have no conflict of interest.

## Extended Data

**Fig. EV1| A.** *c-myc* copy number relative to a reference amplicon on the *Ncl* gene at different time points after LPS stimulation in *c-myc*^*wt/wt*^ and c*-myc*^Δ/Δ^ cells, as indicated. Data are presented as mean ± SD; n = 6 for the t0 and 24h samples, n=4 for the 48h and 72h samples. **B.** *c-myc* mRNA expression (normalized to *TBP*) at different time points after LPS stimulation in *c-myc*^*wt/wt*^ and c*-myc*^Δ/Δ^ cells. n = 3. **C.** as in B, for Myc protein levels, based on quantification of independent immunoblots (n = 4): a representative blot is shown above the plot. Note that in wild-type cells, *c-myc* mRNA levels peaked 2 hours after LPS stimulation(Kelly et al., 1983) while the protein steadily accumulated over time, consistent with post-transcriptional regulation of its synthesis and/or stability (Ehninger, Boch et al., 2014, Farrell & Sears, 2014): as expected, both mRNA and protein accumulation were blunted in c*-myc*^Δ/Δ^ cells. **D.** Myc and IgG ChIP, as indicated, in non-treated (NT) or LPS-treated (2h) *c-myc*^*wt/wt*^ and c*-myc*^Δ/Δ^ B-cells. PCR primers in the *AchR* promoter (as a non-bound control) and in *Ncl* intron 1 (as a known Myc target) were used for quantification (% if input) as previously described (Frank, Schroeder et al., 2001, Guccione et al., 2006). n = 3. **E.** *pus7* and *smyd2* mRNA expression relative to *TBP* at different time points after LPS stimulation in *c-myc*^*wt/wt*^ and c*-myc*^Δ/Δ^ cells. n = 3. **F.** Quantification of total RNA levels per cell along the LPS time-course in *c-myc*^*wt/wt*^ and c*-myc*^Δ/Δ^ cells. n = 3. **G.** Percentages of BrdU positive cells at 0, 12 and 24h, as indicated. n = 3. **H.** Growth curve of *c-myc*^*wt/wt*^ and c*-myc*^Δ/Δ^ cells, as indicated. n = 3. **I.** *c-myc*^*wt/wt*^ and c*-myc*^Δ/Δ^ cells were sorted according to their size and shape (FSC and SSC, respectively) after 24 or 48h of LPS stimulation: cells with low and high FSC+SSC were defined as resting and activated, respectively, and were purified by FACS sorting prior to qPCR quantification of c*-myc* copy number, alongside unsorted c*-myc*^Δ/Δ^ and *c-myc*^*wt/wt*^ control samples. The experiment was repeated twice with similar results. In all the bar plots (except panel I), data are presented as mean ± SD.

**Fig. EV2| A.** Numbers of genes classified in the different regulatory categories on the basis of the RNA-seq data at each time point after LPS stimulation. **B.** Venn diagrams representing the overlap between genes identified as Myc-dependent induced (left) or repressed (right) at the different time points after LPS treatment. **C.** As in B, for Myc-independent LPS-regulated genes.

**Fig. EV3| A.** Heatmap showing the variations in RNAPII ChIP-seq read density (log2FC) in different regions (TSS, gene body, TES) for Myc-dependent genes along the LPS time course, as indicated. The grey scale represents the starting level for each parameter in unstimulated cells. Columns on the left show the changes in Myc share and RNA synthesis, as indicated, rates for the same genes. **B.** As in A, for Myc-independent genes. **C.** Boxplots reporting LPS-induced changes (Log2FC) for each of the four RNAPII kinetic rates in *c-myc*^*wt/wt*^ (wt) and c*-myc*^Δ/Δ^ cells for Myc-dependent induced and repressed genes, as indicated. **D.** As in C, for Myc-independent genes.

**Fig. EV4|** Zoom-in heatmaps and boxplot representations for the clusters represented in Fig. 4D.

**Fig. EV5|** Zoom-in heatmaps and boxplot representations for the clusters represented in Fig. 4E.

**Fig. EV6|** Bar-plots showing the –log10 (FDR) of the top 12 cis-regulatory motifs enriched in the promoters of the genes belonging to the indicated regulatory groups.

**Fig. EV7| A.** Venn diagrams representing the overlap between Myc-dependent LPS-induced genes defined in this study (either all genes, or only those in CL1) with other lists of Myc-regulated genes. From left to right, and top to bottom: genes less expressed in Myc/Max KO B-cells activated with LPS for 72h, compared to wt cells (Perez-Olivares et al., 2018); genes more expressed in Myc-positive versus negative germinal center B-cells (Dominguez-Sola et al., 2012) Myc-dependent serum-responsive (MDSR) genes in fibroblasts (Perna et al., 2012); genes down-regulated in Myc-depleted U2OS (Lorenzin et al., 2016), K562 and HCT116 cells (Muhar et al., 2018). **B.** Venn diagrams representing the overlap between Myc-dependent LPS-induced genes defined in this study with genes induced in Eµ-*myc* B-cells at the pre-tumoral (left) or tumoral stage (right). The p-value of the overlap between Myc-dependent LPS-induced genes and the dataset of interest is shown below each diagram

